# Correlation of IL-17 level in gingival crevicular fluid of orthodontically induced inflammatory root resorption

**DOI:** 10.1101/627059

**Authors:** Hua Zhou, Xiao Liang, Aipeng Liu, Dongmei Nong, Yaqing Qin, Lianxiang Chen, Na Kang

## Abstract

**Objective:** To investigate IL-17 expression in orthodontic tooth movement and orthodontic nickel-titanium spring-induced inflammatory root resorption.

**Methods:** Orthodontic nickel-titanium springs were ligated between the bilateral maxillary first molar and the incisors of the rats to establish a rat model of orthodontic tooth movement (OTM), each rat was subjected to two cycles of near-GCF and peripheral blood serum collection before and after force application, and IL-17 levels in GCF and serum were measured quantitatively by ELISA. Morphological changes in periodontal tissue and root of the experimental dentine were evaluated by hematoxylin and eosin staining. Tartrate-resistant acid phosphatase staining and immunohistochemistry were used to determine the osteoclast number and expression changes in IL-17, receptor activator of nuclear factor kappa-B ligand (RANKL), and osteoprotegerin (OPG) in the periodontal tissues, respectively, on the pressure side of the experimental tooth.

**Results:** IL-17 was detected in GCF and serum. The pressure area exhibited alveolar bone resorption only at a force of 20g. Additionally, a force of 60g led to root resorption. IL-17, RANKL/OPG and osteoclast number showed similar trend that all expressed increasing high level at early stage, then significantly decreased from days 5 to 14, and revealed 60g group the highest expression level while 0g group the lowest.

The change in the IL-17 level in the GCF was strongly correlated with IL-17 and RANKL/OPG expression levels and osteoclast numbers in the periodontal ligament.

**Conclusions:** The results indicated that measuring IL-17 level in GCF can predict the risk of alveolar bone and root resorption induced by orthodontic treatment.

## Introduction

Interleukin-17 (IL-17) is secreted by T-helper 17 (Th17) cells. IL-17 can activate neutrophils to mediate inflammation,^1^ participate in bone resorption in periodontitis,^2^ induce the differentiation of monocyte osteoclasts (OCs),^3^ and induce osteoblasts (OBs) to express receptor activator of nuclear factor κB ligand (RANKL), which generates osteoclasts through the osteoprotegerin (OPG)/RANKL/RANK signal transduction pathway.^4^ Orthodontic treatment can induce root resorption, which is characterized by inflammation and termed orthodontically induced inflammatory root resorption (OIIRR).^5^

Animal studies have demonstrated increased IL-17 expression in gingival crevicular fluid (GCF) on the pressured side.^6^ The positive expression of IL-17 is highly correlated with the relative root resorption area.^6,7,8^ GCF measurements can help confirm the reconstructed state of the periodontium following tooth removal. Enzyme-linked immunosorbent assay (ELISA) to detect IL-17 in GCF and peripheral blood may be useful to confirm that the IL-17 level in peripheral blood correlates with that in GCF.^9,10^

We hypothesized that the force activation of IL-17 production in GCF may contribute to OIIRR. To test this hypothesis, we determined whether orthodontic force induced the activation of IL-17 production in GCF and peripheral blood, and consequently promoted OIIRR by enhancing osteoclastogenesis in a rat model of orthodontic tooth movement (OTM). We investigated whether the IL-17 level in GCF could be indicative of OIIRR.

## MATERIAL AND METHODS

### Animals and Treatments

Sixty-five 8–10-week-old male Wistar rats were used in this study. The experimental protocol was reviewed and approved by the Institutional Animal Care and Use Committee of the university where the study was performed.

### Force Application

The rats were divided randomly into four groups (0g, 20g, 60g, and control). Twenty rats were included in each of the three force groups, and five rats were included in the control group (Table 1). To establish the OTM animal model, orthodontic nickel-titanium coiled springs 1 mm in length (AOJIE, Hangzhou, China) were ligated between the bilateral maxillary first molar and the incisors of the rats using flowable restorative resin to deliver a force of 0g, 20g, or 60g.

The OTM rats were evaluated at different time points (Table 1) to simulate different OIIRR states. Before and after the forces were applied, GCF was collected from the mesio-bilateral maxillary first molar using absorbent paper and orbital venous plexus blood was drawn for use as peripheral blood.

### Tissue Specimen Preparation

The animals were euthanized under deep anesthesia. We selected the intact bilateral maxillary first molar and periodontium as tissue specimens. They were fixed in 4% paraformaldehyde for 24 h (Bio Basic, Markham, Ontario, Canada), decalcified in 10% EDTA-disodium salt for 6–8 weeks (Kelong, Chengdu, China), dehydrated in an alcohol gradient, and embedded in paraffin. Serial parasagittal sections (3–5 μm) were obtained and mounted on glass slides.

### GCF Sampling

GCF samples from the mesial side of bilateral maxillary first molar were collected before and after force application. The study sites were gently dried using an air syringe and were isolated by cotton rolls. Filter paper strips (GAPADENT CO., LTD) were placed gently into the gingival sulcus until a minimum of resistance was felt, left there for 30 s, then put strips into sterilized EP tube. Repeat for eight times at the same site with 1 min interval for each collection. Strips visibly contaminated with blood were discarded. 150 μL phosphate-buffered saline (PBS) 0.05 % (w/v)-Tween-20 buffer was added to each EP tube containing strips and stored −80 °C until the laboratory analyses.

### Serum Sampling

Blood (1 mL) was obtained from the orbital venous plexus and serum was separated by centrifugation (3000 rpm, 15 min, cool). Separated serum samples were collected into microcentrifuge tubes and stored at −80 °C.

### Enzyme-linked immunosorbent assays

ELISA were used for the quantitative detection of IL-17A (eBioscience, America) level in GCF and serum according to published method.^9,10^. Dilution factors were multiplied by the concentration read from a standard curve, generated by plotting the MOD of each standard on the vertical axis versus the corresponding IL-17 standard concentration on the horizontal axis.

### H&E and TRAP Staining

To observe the changes of the root, alveolar bone and periodontal ligament, H&E staining were conducted following the manufacture instruction, images were taken from the pressure side of first molar mesial buccal root with a light microscope(200x). For osteoclast count examination, TRAP staining was performed with a commercial kit (Sigma-Aldrich), randomly selected 5 fields of view on the pressure side of the mesial buccal root of maxillary orthodontic tooth in rats, then the number of osteoclast per bone surface was counted via image-pro-plus6.0.

### Immunohistochemistry

To detect IL-17, RANKL and OPG of periodontal tissue, immunohistochemistry was conducted, specimens were stained respectively with rabbit anti-rat IL-17 primary antibody(1:150, bs-1183R, MBL BeijingBiotech Co., LTD), rabbit anti-rat RANKL primary antibody(1:200, BA1323, BURDOCK Biotech Co., LTD) and rabbit anti-rat OPG primary antibody(1:50, BA1475-1, BURDOCK Biotech Co., LTD), followed by a horseradish peroxidase-conjugated goat anti-rabbit secondary antibody (Zsbio Commerce Store). Calculation of Mean Optical Density (MOD) over randomly selected tension and pressure side of the orthodontic tooth was based on 5 fields per rat, via image-pro-plus6.0 software. A MOD value of 450 nm was set, and MOD value owned a positive correlation with the positive expression of the relative protein.

### Statistical Analyses

Data were analyzed using *t*-tests, one-way analysis of variance, or least significant difference *t*-tests. Spearman rank correlation analysis was performed. A p-value < 0.05 was considered significant.

## RESULTS

### GCF Findings

#### Force-mediated changes in GCF

In general, changes in GCF quantity were significantly different between the 0g, 20g, and 60g groups at all time points except at 1 d (p < 0.05). The changes in GCF levels were significantly different between the 0g and 20g groups at 3 and 5 d (p < 0.05), and between the 0g and 60g groups at 3, 5, 7, and 14 d (p < 0.05). The differences between the 20g and 60g groups reached statistical significance at 5, 7, and 14 d (p < 0.05) (Fig. 1).

#### Changes in IL-17 levels in GCF after force application

In general, changes in the GCF IL-17 levels were significantly different between the 0g, 20g, and 60g groups at all time points except at 1 d (p < 0.05). Changes in the GCF IL-17 levels were significantly different between the 0g and 20g groups at 3, 5, and 7 d (p < 0.05), and between the 0g and 60g groups at 3, 5, 7, and 14 d (p < 0.05). The differences between the 20g and 60g groups reached statistical significance at 5, 7, and 14 d (p < 0.05) (Fig. 2).

### Detection of IL-17 in the Peripheral Blood

The changes in IL-17 levels in peripheral blood were consistent with those for IL-17 levels in GCF (Fig. 3). Changes in the peripheral blood IL-17 levels in the 20g and 60g groups were significantly different than those in the 0 group at 5 and 7 d (p < 0.05), and between the 20g and 60g groups at 7 d (p < 0.05) (Fig. 3).

### H&E Staining Results

#### Non-force pressure

The surfaces of the tooth and alveolar bone in the control and 0g groups were smooth, and osteoclasts were not detected (Fig. 4a and b).

#### 20g pressure

At 3 d, the periodontal ligament began to exhibit bone resorption lacunae and osteoclasts on the alveolar bone surface. At 5 and 7 d, the bone resorption lacunae and osteoclasts were gradually reduced. At 14 d, osteoblasts appeared with a smooth root surface and osteogenic activity (Fig. 4c–g).

#### 60g pressure

At 1 to 3 d, the area of bone resorption lacuna and osteoclasts gradually increased and were mainly located at the bottom third of the root tip. At 5 d, there was a large area of homogeneous parenchyma in the periodontal ligament, and severe bone resorption occurred on the surface of the root and alveolar bone. Many mature osteoclasts were clustered. At 7 d, the range of vitreous changes in the organization, number of osteoclasts, and bone absorption lacunae began to decrease. At 14 d, repair of the root and alveolar bone had begun (Fig. 4h–l).

### IHC Staining Results

#### IL-17 expression

The 0g group displayed weak positive IL-17 staining in individual cells; in contrast, there was a strong positive expression of IL-17 in the 20g and 60g groups in the cytoplasm of fibroblasts and osteoblasts (Fig. 5a–p).

Statistical analysis of the IL-17 MOD values revealed that IL-17 protein expression in the 20g and 60g groups were significantly different than that of the 0g group at each time point (p < 0.05).

The MOD values of IL-17 were significantly different between the 60g and 20g groups at 5 and 7 d (p < 0.05). There were no significant changes in MOD values throughout the study period in the 0g group (Fig. 5q).

#### RANKL expression

On the pressure side, RANKL was positively expressed in the cytoplasm of fibroblasts, osteoblasts, and osteoclast (Fig. 6). In all force groups, the expression trend of RANKL was consistent with that of IL-17 at each time point (Figs. 6b–p).

Statistical analysis of the RANKL MOD values revealed that in the 0g force group, RANKL protein expression did not significantly change. The RANKL MOD values of the 20g and 60g force groups were significantly different than those of the 0g group at each time point (p < 0.05). The RANKL MOD values were significantly different between the 60g and 20g groups at 5, 7, and 14 d (p < 0.05) (Fig. 6q).

#### OPG expression

OPG was positively expressed in the cytoplasm and nucleus of fibroblasts and osteoblasts at all time points in the 0g force group. In the 20g and 60g groups, the expression of OPG showed a tendency to gradually decrease and then increase (Fig. 7a–p).

Statistical analysis of the OPG MOD values revealed that OPG expression in periodontal tissues was not significantly changed in the 0g group. Comparison of the OPG MOD values between the 0, 20, and 60g groups revealed that OPG expression was generally not significantly different at all time points (p < 0.05), except for the 20g group on the first day. The MOD values of OPG were significantly different between the 60g and the 20g groups at 1 and 5 d (p < 0.05) (Fig. 7q).

### TRAP Staining Results

For the TRAP staining, a PBS negative control group was included to show the background staining (Fig. 8a). Osteoclasts were observed as individual cells in the 0g group. In the 20g and 60g groups, the number of osteoclasts increased at 1 d, peaked at 3 and 5 d, respectively, and then decreased gradually (Fig. 8b–p).

Statistical analysis of the osteoclast count revealed that the number of osteoclasts in the 0g group did not significantly change. Generally, the osteoclast numbers in the 20g and 60g groups were significantly different than those in the 0g group at all time points (p < 0.05), except at 1 d for the 20g group. Osteoclast numbers were significantly different between the 60g and the 20g groups at 5, 7, and 14 d (p < 0.05) (Fig. 8q).

### Correlation Analysis Results

Spearman rank correlation analysis revealed that the level of IL-17 in GCF was moderately correlated with the level of IL-17 in peripheral blood and was highly correlated with the MOD value of IL-17 in periodontal tissue, the ratio of RANKL/OPG, and the number of osteoclasts (p < 0.05). The MOD value of IL-17 in periodontal tissue was highly correlated with the ratio of RANKL/OPG and the number of osteoclasts (p < 0.05). The ratio of RANKL/OPG in periodontal tissue was significantly correlated with the number of osteoclasts (p < 0.05) (Table 2).

## DISCUSSION

This study aimed to determine whether IL-17, RANKL, and OPG are involved in root resorption during orthodontic treatment by applying excessive orthodontic forces in a rat model. Under the action of orthodontic force, the vascular activity in the periodontal ligament increases, which leads to changes in cell morphology that results in vasodilatation and exudation of leukocytes to the extravascular space, and the release of cell factors, histolytic enzymes, and acidic products. These events eventually lead to increased GCF flow,^11^ whereas changes in GCF flow occur earlier than changes in its composition.^12^

At 3 d, the levels of GCF and IL-17 increased gradually in the 0g group and then reached a stable level. The gradual increase indicates that the OTM device affected the self-cleaning of teeth by the rats, causing mild gingival inflammation. However, as indicated by the stabilization of the GCF and IL-17 levels, the inflammation did not intensify. The levels of GCF and IL-17 in the GCF of rats in the 0g group were significantly different from those in the 20g and 60g groups. These results indicate that the orthodontic force had a certain effect on the level of GCF and IL-17 in GCF when excluding periodontal stimulation. Thus, the levels of GCF and IL-17 in GCF after force application were influenced by the gingival inflammation and the alteration of the deep periodontal tissue. The concentration of IL-17 in GCF was increased when the force increased.

The level of IL-17 in GCF after orthodontic treatment has been measured in clinical studies. IL-17 in GCF at the orthodontic-treated tooth was reportedly higher than that before treatment and on the tension side.^7^ The concentration of IL-17 was shown to correlate with the magnitude of the orthodontic force.^8^ When the level of IL-17 in GCF changes, the relevant components in peripheral blood also change.^13^

The levels of IL-17 in GCF and peripheral blood were affected by inflammatory factors on the gingival surface and the remodeling of deep periodontal tissues. Changes in GCF levels of IL-17 and serum levels of IL-17 were moderately correlated, indicating that there is a correlation between peripheral blood- and GCF-related factors and suggesting that peripheral blood CD4^+^ Th17 cells play a key regulatory role in orthodontic processes via the IL-17 pathway.

IL-17 can directly stimulate osteoclastogenesis or promote osteoclast formation by the up-regulation of osteoclast differentiation-related factors and plays a role in promoting bone resorption.^14^ There are many factors and pathways involved in the up-regulation of osteoclasts during bone resorption. The OPG/RANKL/RANK pathway is considered the key protein loop that regulates osteoclast differentiation and activation, with RANKL being a key factor in osteoclast differentiation and activation and inhibition of osteoclast apoptosis.^15^ In a rat model of periapical periodontitis, the expression of IL-17 and RANKL mRNA indicated that IL-17 induces osteoclast formation by regulating the expression of RANKL, resulting in bone resorption.^16^

Under 20g of force, the expression of IL-17 and RANKL peaked at 3 d, alveolar bone resorption was active, and no absorption was observed on the root surface. Under 60g of force, alveolar bone absorption and lacunae on the root surface began to appear. The trend for MOD value of IL-17 by IHC was similar to that of RANKL MOD value. The latter MOD value also displayed a similar trend with that of the osteoclast number, consistent with our hypothesis that IL-17 can promote osteoclast differentiation and maturation by up-regulating RANKL, which is involved in root and alveolar bone absorption.

The combination of high levels of OPG and RANK can inhibit RANKL involvement in osteoblast differentiation. The RANKL/OPG system is involved in the formation of osteoclasts and regulates the process of root resorption.^17^ In the present study, the MOD value of OPG displayed an opposite trend than the MOD value of IL-17 and RANKL and the number of osteoclasts. The ratio of RANKL/OPG was significantly correlated with the MOD value of IL-17 and the number of osteoclasts. Therefore, we speculate that under the action of orthodontics, the levels of IL-17 and RANKL increased on the pressure side of the tooth. The increase in RANKL resulted in the competitive reduction of OPG, followed by an increase in the RANKL/OPG ratio, promoting the increase in osteoclasts and the subsequent alveolar bone and root absorption.

The detection of IL-17 levels in GCF may be used to quickly assess orthodontic force in a chair-side operation.^8^ In this study, the change in the IL-17 level in GCF after orthodontic force was applied was positively correlated with the MOD value of IL-17. In addition, the change in the IL-17 level in GCF after force application was significantly correlated with the ratio of RANKL/OPG and the number of osteoclasts in periodontal tissues, indicating that the changes in the IL-17 level in GCF can predict changes in the IL-17 level in deep periodontal tissue. The changes in the IL-17 level in GCF in the 20g and 60g groups at 5, 7, and 14 d were significantly different (p < 0.05). Thus, the expression of IL-17 in GCF in the root resorption group was higher than that in the non-absorption group, indicating that root resorption may have occurred when an excess of IL-17 in GCF was detected.

## CONCLUSION

The changes in IL-17 in deep periodontal tissues can be detected in GCF. Measuring the level of IL-17 in GCF can effectively predict the change in IL-17 in periodontal tissues and be used to predict the extent of alveolar bone and root absorption under orthodontic treatment. Orthodontic root resorption has always been the focus of attention. Collection of gingival crevicular fluid in a clinical setting is a painless process, which is more easily accepted by patients, and it can also help to monitor root resorption more effectively and quickly, leading to further standardization of clinical procedures.

## Abbreviation

OTM: Orthodontic Tooth Movement
Interleukin-17: IL-17
GCF: Gingival Crevicular Fluid
RANKL: Receptor Activator of Nuclear factor Kappa-Β Ligand

## Acknowledgements

This study was supported by the Guangxi medical university animal experimental center, Naning, China.

## Availability of data and materials

All data available within the manuscript in the form of results, tables, and figure.

## Ethics approval

The animal experimental protocol in this study was approved by the Ethics Committee for Animal Experiments in Guang Xi Medical University with the number GXMU/Animal Ethics Approval/20130308-14.

## Consent for publication

Authors have given their consent for their data to be published in the report.

## FIGURE LEGENDS

Fig. 1 Changes in gingival crevicular fluid (GCF) levels in the different experimental groups. Orthodontic nickel-titanium coiled springs were used to deliver a force of 0g, 20g, or 60g, and GCF was collected at the indicated time points using absorbent paper. Data represent the mean differences in GCF quantity (vs. control rat) ± SD; *n* = 4 per experimental group per time point; *, p < 0.05 between the 0g and 20g or 60g groups; Δ, p < 0.05 between the 20g and 60g groups.

Fig. 2 Changes in IL-17 levels in gingival crevicular fluid (GCF) in the different experimental groups. The level of IL-17 in the GCF of rats in the 0g, 20g, and 60g groups was assessed by ELISA, and data are presented as changes in the GCF IL-17 levels (vs. control rat) ± SD; *n* = 4 per experimental group per time point; *, p < 0.05 between the 0g and 20g or 60g groups; Δ, p < 0.05 between the 20g and 60g groups.

Fig. 3 Changes in serum IL-17 levels in the different experimental groups. The level of IL-17 in the peripheral blood of rats in the 0g, 20g, and 60g groups was assessed by ELISA, and data are presented as changes in the serum IL-17 levels (vs. control rats) ± SD; *n* = 4 per experimental group per time point; *, p < 0.05 between the 0g and 20g or 60g groups; Δ, p < 0.05 between the 20g and 60g groups.

Fig. 4 Hematoxylin and eosin (HE) staining of intact bilateral maxillary first molar and periodontium. Representative HE images (×200) from the control, 0g, 20g, and 60g groups are shown; n = 1 for the control; *n* = 4 for each experimental group per time point. Scale bars = 50 μm; R = root; black solid arrows = bone resorption pits; black dashed arrows = osteoclasts; red solid arrows = homogeneity of hyaline tissue.

Fig. 5 IL-17 protein expression in periodontal tissue by immunohistochemistry (IHC). (a) Negative staining and (b–p) representative IL-17 IHC images (×400) from the 0g, 20g, and 60g groups are shown; *n* = 4 per experimental group per time point; scale bars = 20 μm; R = root. (q) Mean optical density (MOD) of IL-17 IHC; *, p < 0.05 between the 0g and 20g or 60g groups; Δ, p < 0.05 between the 20g and 60g groups. In the 0g group, weak-positive IL-17 expression was observed from 1 to 14 d. In the 20g group, IL-17 expression gradually increased at 1 d, peaked at 3 d, and gradually decreased at 5, 7, and 14 d. In the 60g group, IL-17 gradually increased at 1 d, peaked at 5 d, and gradually decreased at 7 and 14 d.

Fig. 6 RANKL protein expression in periodontal tissue by immunohistochemistry (IHC). (a) Negative staining and (b–p) representative RANKL IHC images (×400) from the 0g, 20g, and 60g groups are shown; *n* = 4 per experimental group per time point; scale bars = 20 μm; R = root. (q) Mean optical density (MOD) of RANKL IHC; *, p < 0.05 between the 0g and 20g or 60g groups; Δ, p < 0.05 between the 20g and 60g groups. In the 0g group, weak-positive RANKL expression was observed at 1 to 14 d. In the 20g group, the positive expression of RANKL gradually increased at 1 d, peaked at 3 d, and then gradually decreased at 5, 7, and 14 d. In the 60g group, RANKL expression gradually increased at 1, peaked at 5, and gradually decreased at 7 and 14 d.

Fig. 7 OPG protein expression in periodontal tissue by immunohistochemistry (IHC). (a) Negative staining and (b–p) representative OPG IHC images (×400) from the 0g, 20g, and 60g groups are shown; *n* = 4 per experimental group per time point; scale bars = 20 μm; R = root. (q) Mean optical density (MOD) of OPG IHC; *, p < 0.05 between the 0g and 20g or 60g groups; Δ, p < 0.05 between the 20g and 60g groups. In the 0g group, from 1 to 14 d, positive expression of OPG was observed. In the 20g group, OPG expression gradually decreased from 1 d, reached the lowest level at 3 d, and increased gradually at 5, 7, and 14 d. In the 60g group, OPG expression gradually decreased from 1 d, reached the lowest level at 5 d, and increased gradually at 7 and 14 d.

Fig. 8 Tartrate-resistant acid phosphatase (TRAP) staining to observe mature osteoclasts in periodontal tissue. (a) Negative staining (PBS only) and (b–p) representative TRAP staining images (×400) from the 0g, 20g, and 60g groups are shown; *n* = 4 per experimental group per time point; scale bars = 20 μm; R = root. (q) Osteoclast count based on the TRAP staining results; *, p < 0.05 between the 0g and 20g or 60g groups; Δ, p < 0.05 between the 20g and 60g groups.

## REFERENCES

[1] Manel N, Unutmaz D, Littman DR. The differentiation of human T(H)-17 cells requires transforming growth factor-beta and induction of the nuclear receptor RORγt. Nat Immunol 2008;9:641–9.

[2] Trombone AP, Ferreira SB Jr, Raimundo FM, de Moura KC, Avila-Campos MJ, Silva JS, et al. Experimental periodontitis in mice selected for maximal or minimal inflammatory reactions: increased inflammatory immune responsiveness drives increased alveolar bone loss without enhancing the control of periodontal infection. J Periodontal Res 2009;44:443–51.

[3] Yago T, Nanke Y, Ichikawa N, Kobashigawa T, Mogi M, Kamatani N, et al. IL-17 induces osteoclastogenesis from human monocytes alone in theabsence of osteoblasts, which is potently inhibited by anti-TNF-α antibody: a novel mechanism of osteoclastogenesis by IL-17. J Cell Biochem 2009;108:947–55.

[4] Zhang F, Wang CL, Koyama Y, Mitsui N, Shionome C, Sanuki R, et al. Compressive force stimulates the gene expression of IL-17s and their receptor in MC3T3-E1 cells. Connect Tissue Res 2010;51:359–69.

[5] Brezniak N, Wasserstein A. Orthodontically induced inflammatory root resorption. Part I: the basic science aspects. Angle Orthodo 2002;72:175–9.

[6] Zhang J-P, Gao H, Xiao D-N, Liu D-Y. Interleukin-17 levels in gingival crevicular fluid of orthodontic patients. Beijing Journal of Stomatology 2011;19:144–146.

[7] Song N, Jin S-M, Ma Y-W, Xu J-G, Zhang J. The study of change of the amount of GCF before and after orthodontic tooth movement of rats. Journal of Clinical Stomatology 2009;25:209–211.

[8] Fan R-X, Li X-Z. Interleukin-17 levels in gingival crevicular fluid of orthodontic tooth. IMMUNOLOGICAL JOURNAL 2014;30:906–909.

[9] Awang RA, Lappin DF, MacPherson A, Riggio M, Robertson D, Hodge P, et al. Clinical associations between IL-17 family cytokines and periodontitis and potential differential roles for IL-17A and IL-17E in periodontal immunity. Inflamm Res 2014;63:1001–12.

[10] Cifcibasi E, Koyuncuoglu C, Ciblak M, Badur S, Kasali K, Firatli E, et al. Evaluation of local and systemic levels of interleukin-17, interleukin-23, and myeloperoxidase in response to periodontal therapy in patients with generalized aggressive periodontitis. Inflammation 2015;38:1959–68.

[11] Davidovitch Z, Nicolay OF, Ngan PW, Shanfeld JL. Neurotransmitters, cytokines, and the control of alveolar bone remodeling in orthodontics. Dent Clin North Am 1998;32:411–35.

[12] Baldwin PD, Pender N, Last KS. Effects on tooth movement of force delivery from nickel-titanium archwires. Eur J Orthod 1999;21:481–9.

[13] Zhou Y, Xu Y. Changes of Th17 cytokines and RORC in chronic periodontitis patients. Chinese Journal of Conservative Dentistry 2012;22:103–106.

[14] Lubbers E, Koenders MI, Vandenberg WB. The role of T-cell interleukin-17 in conducting destructive arthritis: lessons from animal model. Arthritis Res Ther 2005;7:29–37.

[15] Udagawa N, Takahashi N, Jimi E, et al. Osteoblasts/stromal cells stimulate osteoclast activation through expression of osteoclast differentiation factor/RANKL but not macrophage colony-stimulating factor:receptor activator of NF-kappa B ligand. Bone 1999;25:517–23.

[16] Wei S-S, Kawashima Nobuyuki, Suzuki Noriyuki, Xu J, Sudahi deaki. Expression of Th17-related cytokines in induced rat periapical lesions. Beijing Journal of Stomatology 2010;18:21–24.

[17] Michigami T, Ihara W, Yamazaki M, Ozono K. Receptor activator of nuclear factor kappaB ligand (RANKL) is a keymolecule of osteoclast formation for bone metastasis in a newly developed model of human neuroblastoma. Cancer Res 2001;61:1637–44.

